# Coinfection with malaria alters the dynamics and fitness of an intestinal nematode

**DOI:** 10.1101/2025.09.15.676228

**Authors:** Luc Bourbon, Aloïs Dusuel, Emma Groetz, Mickaël Rialland, Benjamin Roche, Bruno Faivre, Gabriele Sorci

## Abstract

Infections with soil transmitted helminths (STHs) are highly prevalent in humans living in the intertropical region. While, in most cases, STHs can establish chronic infections, the dynamics of the infection can be altered when other parasites exploit the same host. These changes can have consequences in terms of the health of the host, the epidemiology of the disease (e.g., the duration of the infection and the inter-host transmission success) and the fitness of the parasite. Here, we investigated if the coinfection with *Plasmodium yoelii* alters the dynamics and the fitness of the murine nematode *Heligmosomoides polygyrus*. We found that, compared to single infected mice, coinfection produced an increase in the number of excreted eggs, while the biomass of adult worms in the intestine did not differ between single infected and coinfected mice. Moreover, the increase in egg excretion was also observed when *Plasmodium* infected hosts that had been harboring the nematode during the past four weeks (i.e., when the population size of adult worms can only decrease due to mortality). Therefore, the enhanced shedding of eggs reflects a plastic adjustment of worm fecundity to the environment provided by a coinfected host. This plastic response was modulated by the host Th2 immunity, as coinfection inhibited *IL-4* and *IL-13* gene expression, plasma levels of IL-5 and IL-13, and the expansion of GATA-3^+^ CD4^+^ T cells in the spleen. In agreement with this, experimentally inhibiting IL-13 with neutralizing antibodies reproduced the results observed in coinfected mice (an increase in egg excretion), while the administration of recombinant IL-13 reduced egg shedding. Interestingly, coinfection had a net positive effect on parasite fitness as shown by a longer persistence within the host and higher cumulative number of eggs excreted up to 99 days post-infection. Although the gene expression of Th2 cytokines was lower at day 99 p.i., coinfected mice still had a downregulated expression compared to single infected hosts. These results show that coinfection with *Plasmodium* has the potential to affect the epidemiology of STHs by increasing the number of eggs excreted over the whole infectious period and maintaining a larger environmental reservoir of transmissible stages.

**Author Summary:** Coinfection between soil-transmitted helminths and malaria is common in several countries of the intertropical region, especially among the most vulnerable populations. Coinfection has the potential to worsen the symptoms caused by malaria, therefore it is important to understand what are the epidemiological and ecological factors that promote the occurrence of coinfection. Transmission of soil-transmitted helminths usually requires human contact with transmissible stages (parasitic eggs or larvae) in the environment; therefore, high egg excretion in the feces of infected people is a key factor contributing to maintain a reservoir of infective stages from which humans can get infected. In this study, we experimentally investigated whether coinfection with malaria alters the dynamics (egg excretion, infection persistence) of a murine intestinal nematode. We found that hosts infected with malaria and subsequently infected with the nematode, excreted more nematode eggs for a longer period, compared to single infected hosts. These changes were mediated by an impaired Th2 immune response in coinfected hosts. These results suggest that malaria coinfection produces positive feedback on key epidemiological traits of the nematode that can further enhance the risk of malaria/helminths cooccurrence.

## Introduction

Soil-transmitted helminths (STHs) are agents of highly prevalent parasitic diseases in most countries of the intertropical region, associated with poverty and unfavorable socioeconomic conditions (Brooker 2010, Loukas et al. 2021). According to the World Health Organization, 24% of the world’s population suffers from infection with STHs (c.a. 1.5 billion of humans) (WHO 2023). Infection with STHs includes several species of nematodes that differ in terms of transmission mode, duration and intensity of the infection, or morbidity. STHs are transmitted either through the ingestion of food and/or water that has been contaminated by the eggs of the helminths (roundworms and whipworms) or through the penetration of the host skin by the infective larvae (hookworm). STHs usually establish chronic infections that can last for years (Loukas et al. 2021). Parasite burden is a key driver of the morbidity of the infection (Else et al. 2020) and severe symptoms involve hookworm-induced anemia in children and pregnant women, roundworm-induced intestinal obstruction, tissue damage during larval migration and impaired physical development (Lustigmann et al. 2012, Loukas et al. 2021), although there is still a paucity of data (Campbell et al. 2016, Raj et al. 2022). *Strongyloides stercoralis* can also cause damage due to its unique capacity (among STHs) to shortcut the environmental stage of the life cycle and directly reinfect the host, especially in immune compromised individuals (Tamarozzi et al. 2019). However, for the vast majority of STHs, infection involves the contact with transmissible stages in the external environment; therefore, high egg excretion in the feces of infected people is a key factor contributing to maintain a reservoir of infective stages from which humans can get infected. Mass treatment with anthelmintic drugs can reduce the prevalence of the infection and alleviate the symptoms of the disease (Marocco et al. 2017; Stroffolini et al. 2023), but the consistency of the benefits has been debated (Humphries et al. 2012, Allen and Parker 2016). Moreover, worm resistance to drugs raises concerns over the sustainability of anthelmintic mass treatment on the long-term (Vercruysse et al. 2011, Prichard et al. 2012, Tinkler 2020, Coffeng et al. 2024). This stresses the importance of understanding the environmental factors (including host immunity) that modulate the production of transmissible stages and the associated transmission success and infection risk (Oyesola et al. 2022).

Geographic areas with high prevalence of helminthiases are also endemic for major infectious diseases, such as malaria or AIDS and therefore people can harbor multiple infections involving macro and microparasites (Webb et al. 2012, Degarege et al. 2016, Afolabi et al. 2021, Sumbele et al. 2021). Coinfection between STHs and malaria has the potential to exacerbate the severity of the disease symptoms and substantial effort has been devoted to understand under which circumstances STHs might aggravate *Plasmodium* virulence (Nacher 2011, Kwan et al. 2018, Hürlimann et al. 2019). On the contrary, the effect of coinfection on the dynamics of STHs has been barely addressed, and clearly merits further investigation.

The reasons underlying a possible effect of coinfection on STH dynamics are manifold. In particular, infection with *Plasmodium* (or other microparasites) polarizes the host immune response towards a type 1 immunity (Olatunde et al. 2022), while control of helminth infection requires a type 2 response orchestrated by Th2 cytokines such as IL-4, IL-5 or IL-13 (Rausch et al. 2008, Allen and Maizels 2011, Maizels et al. 2012, Grencis 2015, Mabbott 2018). Therefore, when hosts are infected with microparasites that elicit a Th1 response, they should be less able to resist a helminth infection (Hoeve et al. 2009, Coomes et al. 2015, Ahmed et al. 2017), implying that when exposed to helminth infective stages, the probability of successful infection should be higher in coinfected hosts. Alternatively, previous infection might debilitate the host and limit its access to resources, offering a more favorable ground for subsequent infections, independently from the host capacity to mount any parasite-specific immune response (Budischak et al. 2015).

The outcome of the coinfection might also depend on the order of the infections with microparasites and helminths (Karvonen et al. 2019, Billet et al. 2020). Previous work has shown that hosts previously infected with helminths suffer from more severe disease symptoms when subsequently infected with *Plasmodium* (Salazar-Castañón et al. 2014, 2018, Dusuel et al. 2025), or other intracellular parasites, such as *Toxoplasma gondii* (Szabo et al. 2024). These results are also in agreement with the idea that initial polarization of the immune response towards a Th2 response makes the host more susceptible to *Plasmodium* (Supali et al. 2010). Therefore, it is important to understand whether the predicted changes in host susceptibility to helminths also depend on the order of infection.

Here, we used an experimental model involving two murine parasites [the intestinal nematode *Heligmosomoides polygyrus* (hereafter Hp) and *Plasmodium yoelii* (hereafter Py)] to investigate if coinfection had any effect on the dynamics and fitness of Hp, and if these effects were driven by the host immune response. In addition, we investigated if coinfection has any long-term effect on Hp dynamics, by assessing the persistence of the infection within the host and the cumulative number of eggs excreted.

We predicted that Hp infecting hosts harboring Py should have higher infection success and therefore excrete more eggs in the host feces. However, if coinfected hosts have an impaired immune response towards Hp, accounting for an enhanced egg shedding, the higher investment into reproductive investment should incur a cost for worms in coinfected hosts. Therefore, we also predicted that the positive effect of coinfection on Hp egg excretion should be transitory, resulting in similar fitness in single infected and coinfected hosts.

## Materials and Methods

### Ethical statement

Experiments were approved by the Comité d’éthique Grand Campus Dijon of the University of Burgundy and were authorized by the French Ministry of Higher Education and Research under the numbers 33492 and 47487.

### Experimental animals and infections

C57BL/6JRj female mice (7-10 weeks old) were purchased from Janvier Labs (Le Genest-Saint-Isle, France), housed in cages containing 5 individuals under pathogen-free conditions, and maintained under a constant temperature of 24°C and a photoperiod of 12h:12h light:dark with ad libitum access to water and standard chow diet (A03-10, Safe, Augny, France). All mice were acclimatized to the housing conditions during, at least, 7 days prior to the start of the experiments, were monitored twice a day to check health status, and euthanized by cervical dislocation under anesthesia with isoflurane either if they reached previously defined end points or at specific days (14, 21 and 35) post-infection (p.i.) for terminal collection of blood and organs.

Mice were infected with *Heligmosomoides polygyrus bakeri* by oral gavage with L3 larvae (350 larvae suspended in 0.2 ml of drinking water) and with *Plasmodium yoelii* 17XNL by intra peritoneal injection (i.p.) with 5x10^5^ infected red blood cells (iRBC) suspended in 0.1 ml PBS. To obtain Hp L3 larvae, feces from donor infected mice were mixed with distilled water and charcoal, spread on a Whatmann paper laid on a wet paper towel and placed into a Petri dish. Petri dishes were stored at room temperature in the dark. L3 larva were collected after 11 days by washing out the back side of the Whatmann paper, the paper towel and the bottom of the Petri dish with distilled water. L3 were then isolated by centrifugation (100 rpm, 10 min, 4°C), washed twice, and kept at 4°C in distilled water until use.

### Experimental groups

Mice were randomly assigned to 5 experimental groups (10 to 15 mice per group). A control group of mice was sham infected (i.p. injection of 0.1 ml of PBS and oral gavage with a 0.2 ml of drinking water); one group received a single Hp infection; one group was infected with Py and 14 days after with Hp; one group was infected with Py and 28 days after with Hp; one group was infected with Hp and 28 days after with Py. The timings of infection in the coinfection groups correspond to specific stages in the life cycles of the two parasites. Day 14 and day 28 post Py infection approximately correspond to the parasitemia peak and the end of the acute phase infection, respectively. Day 28 post Hp infection corresponds to the stage when all larvae have molted into adults, adults have emerged into the intestinal lumen and the infection has reached a chronic stage. The whole experiment was repeated twice, although not all traits were measured in all mice of the two replicates. Additionally, ten mice per group (excluding the non-infected group) were monitored up to 99 days post Hp infection to assess the effect of coinfection on Hp persistence (i.e., clearance rate) and the cumulative egg shedding.

### Hp egg shedding and infection burden

To assess the fecal egg count (FEC, number of parasite eggs per gram of feces), mice were individually placed for 1 hour in clean cages; 200 mg of feces were then collected and disposed in tubes containing 2.5 ml of water. Feces were mechanically broken up and 5 ml of salted water (75% saturation, 0.27g NaCl per ml of water) were added to allow egg floatation. Eggs were counted in a McMaster chamber using an optical microscope (Eclipse E400, Nikon, Tokyo, Japan) under magnification x150. To assess infection burden, the intestines were removed, transferred in tubes containing a deep freeze solution (4.2g of sorbitol, 100ml of 0.9% NaCl solution and 39ml of glycerol) and stored at -80° C. Intestines were then dissected and all worms found were collected, washed with water and disposed in 1.5 ml tubes (that had been previously weighed using a precision balance (MCA6.6S-2S00-M, Sartorius, Göttingen, Germany) (± 1µg) with 500 µl of absolute ethanol. The opened tubes were subsequently placed in an incubator at room temperature during 2 – 3 days. When dry, tubes were weighed again and the worm biomass assessed as the difference between the second and first measurement of tube weights.

### Plasma and organ processing

Plasma was separated from total blood by centrifugation (7 min 3000 rpm 4°C), aliquoted and stored at -80°C. About one third of the spleen was cut and immediately frozen in liquid nitrogen for RT-qPCR, and the remaining was kept in cold PBS before the flow cytometry staining procedure.

### IL-13 administration and neutralization

Mice (8 weeks) were infected with Hp and randomly split into three groups at day 28 p.i. : one group (n = 8) received three i.p. injections of 2 µg recombinant mouse IL-13 (rIL-13) (PeproTech® 210-13-50UG, Gibco, Grand Island, New York, USA) in 200 ml of PBS; one group (n = 7) received three i.p. injections of 100 µg anti-mouse IL-13 monoclonal Ab (clone eBio1316H, Invitrogen™, Carlsbad, California, USA) in 200 ml of PBS; one group (n = 7) received three i.p. injections of 100 µg control IgG1 (clone eBRG1, Invitrogen™) in 200 µl of PBS. Mice were injected at days 28, 30 and 32 post-infection.

### RNA extraction and RT-qPCR for IL-4 and IL-13 gene expression

Spleens were homogenized in TRIzol™ Reagent (Invitrogen™, Carlsbad, California, USA) under strong agitation using 0.5 mm glass beads and Precellys® 24 Touch homogenizer (Bertin Technologies, Montigny-le-Bretonneux, France). RNA extraction was performed following the manufacturer’s instructions. RNA concentration was measured with NanoPhotometer® N50 (Implen, Munich, Germany). Reverse transcription was performed with the High-Capacity cDNA Reverse Transcription Kit (Applied Biosystems™, Foster City, California, USA) from 1.5 µg total RNA. Quantitative PCR were performed with PowerUp™ SYBR™ Green Master Mix (Applied Biosystems™) on QuantStudio™ 3 Real-Time PCR System (Applied Biosystems™). We used two housekeeping genes (β-actin and GAPDH); β-actin provided more consistent values (less intra-group variability) and therefore we used it as housekeeping gene. Primer sequences are reported in the supplementary material (table S1).

### Plasma cytokine quantitation

Concentrations of plasma IL-5 and IL-13 were measured by multiplex using a Mouse Luminex® Discovery Assay (LXSAMSM, Bio-Techne, Minneapolis, Minnesota, USA) according to the manufacturer’s instructions. Samples were analyzed with a Bio-Plex 200 system (Bio-Rad, Hercules, California, USA) in the ImaFlow Facility (US58 BioSanD, Dijon, France).

### Spectral flow cytometry

Spleens were homogenized with a 70 µm cell strainer and washed two times with PBS to obtain a single-cell suspension. Red blood cells were removed from suspensions with eBioscience™ RBC Lysis Buffer (Invitrogen™) (3 min incubation at room temperature). Live cells were counted with viability trypan blue dye and dispatched in 96-well V bottom plates to obtain 1.10^7^ cells/well. Cells were then stained with LIVE/DEAD™ Fixable Blue Dead Cell Stain Kit (Invitrogen™) in PBS for 30 min at 4°C, incubated in stain buffer with BD Fc Block™ (BD Pharmingen™, Franklin Lakes, New Jersey, USA) for 15 min at room temperature (0.5 µg/well), and stained with surface antibodies in brilliant stain buffer (see table S2 for details on the markers used) for 30 min at 4°C. Cells were then fixed and permeabilized with eBioscience™ GATA-3 CD4 Transcription Factor Staining Buffer Set (Invitrogen™) prior intracellular staining in brilliant stain buffer (table S2) for 30 min at room temperature. Cells were resuspended in stain buffer, filtered with 100 µm nylon mesh and analyzed on a 4-laser Cytek Aurora™ spectral flow cytometer (Cytek® Biosciences, Fremont, California, USA). Events were gated and analyzed with SpectroFlo® v3.0.0 software (Cytek® Biosciences) (see figure S1 for the gating strategy).

### Statistical analyses

FEC were analyzed using general linear mixed models (GLMM) where mouse ID was included as a random intercept. FEC were log-transformed to reduce skewness and we checked the distribution of the model residuals. Models also included the treatment group and time p.i. as fixed effects. For variables that were measured only once per individual mouse, we ran general linear models (GLM) with the same predictors, except the random effect. The interaction between the treatment group and time was dropped from the models if the p value was above the 0.05 threshold. We ran models where the coinfection groups were either kept separated (and referred to as *Treatment*) or clustered together (and referred to as *Infection*). Post-hoc comparisons between groups were done using LS-means with Bonferroni correction of p values. Differences in sample size across groups reflect missing values (for instance when mice did not defecate during the time they were placed in the individual cages, or due to technical issues during the processing of the samples).

The persistence of the infection was analyzed using the Kaplan Meier estimator and significance assessed using a Log-Rank test. We considered that mice that shed no eggs during at least two consecutive sampling dates had cleared the infection. For the analysis of relative fitness, we added the number of eggs excreted during each of the 14 sampling dates (from day 12 to day 99 p.i.) for each mouse in each experimental group. These sums were then divided by the mean of the sums for the single Hp infection group. These values therefore refer to the relative fitness with respect to single infection.

All the statistical analyses and figures were done using SAS Studio.

## Results

### Effect of coinfection on Hp egg shedding and infection burden

There was no time-dependent effect on egg shedding over the day 11 to day 35 p.i. (table S3, fig. S2a). However, there was a difference in the average FEC between groups with coinfected mice shedding significantly more eggs compared to single Hp infected individuals (table S3, fig. S2a). Clustering the two coinfection treatments in a group and comparing it to the single infection treatment showed that, overall, coinfected mice shed more eggs than single infected hosts (table 1, fig. 1a).

**Figure 1.**
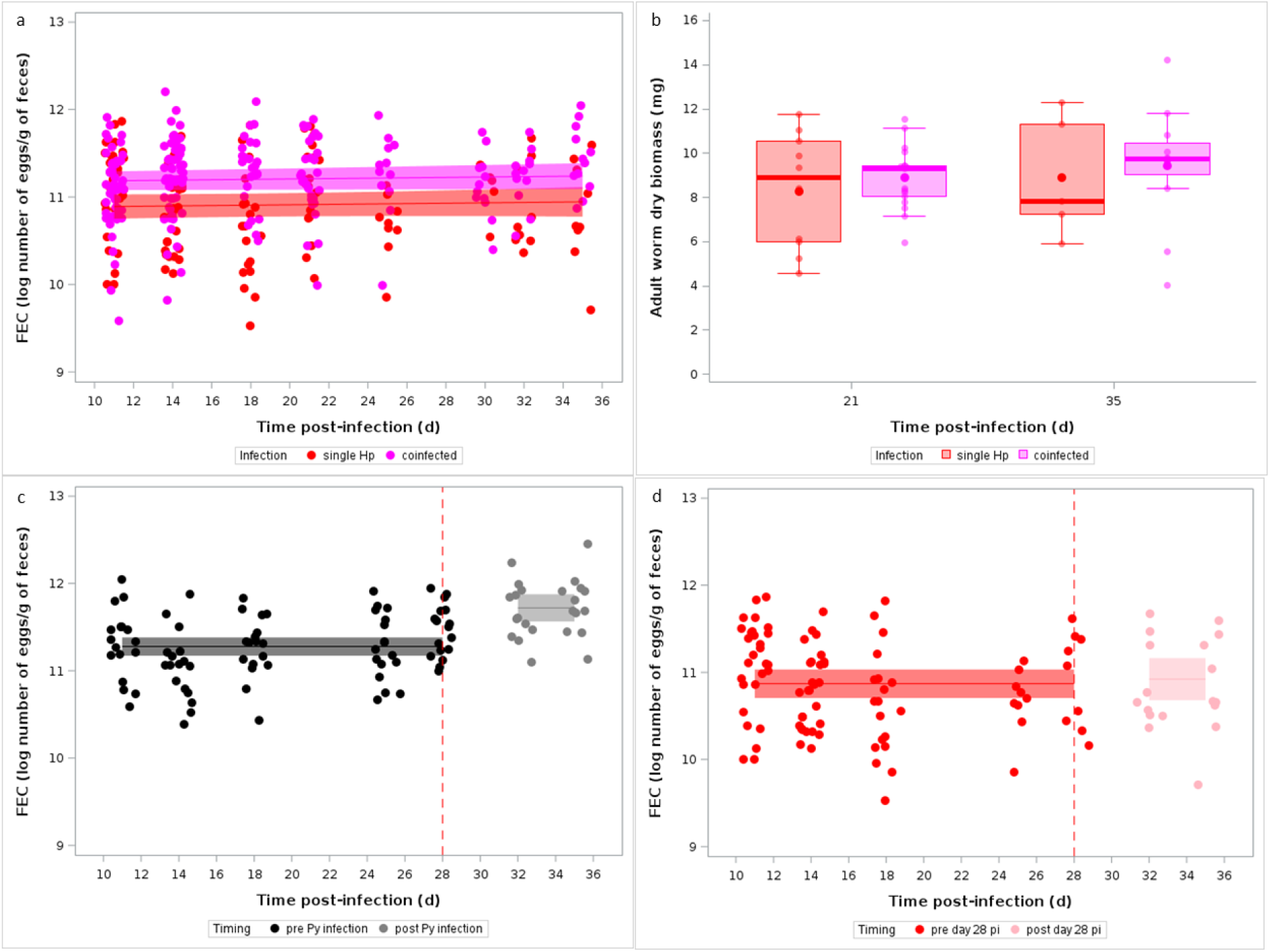
Coinfection with Py improves the fecundity of Hp. a) Fecal egg count (log number of eggs per gram of feces) over the course of the infection for single Hp infected and coinfected (with Py infecting first) mice. Dots represent the raw data, the lines represent the fit of a GLMM, and the shaded area around the line the 95% CI. b) Biomass of adult Hp collected at day 21 and 35 p.i. in the intestinal lumen of single Hp infected and coinfected (with Py infecting first) mice. Dots represent the raw data, the boxes represent the interquartile range (IQR), the horizontal lines the median, and whiskers the range of data within 1.5 the IQR. c) Changes in fecal egg count (log number of eggs per gram of feces) following the infection with Py at day 28 post Hp infection (dotted vertical line). Dots represent the raw data, the lines represent the fit of a GLMM, and the shaded area around the line the 95% CI. d) Changes in fecal egg count (log number of eggs per gram of feces) up to day 28 and after day 28 p.i. (dotted vertical line) in single Hp infected mice. Dots represent the raw data, the lines represent the fit of a GLMM, and the shaded area around the line the 95% CI.

**Table 1.**
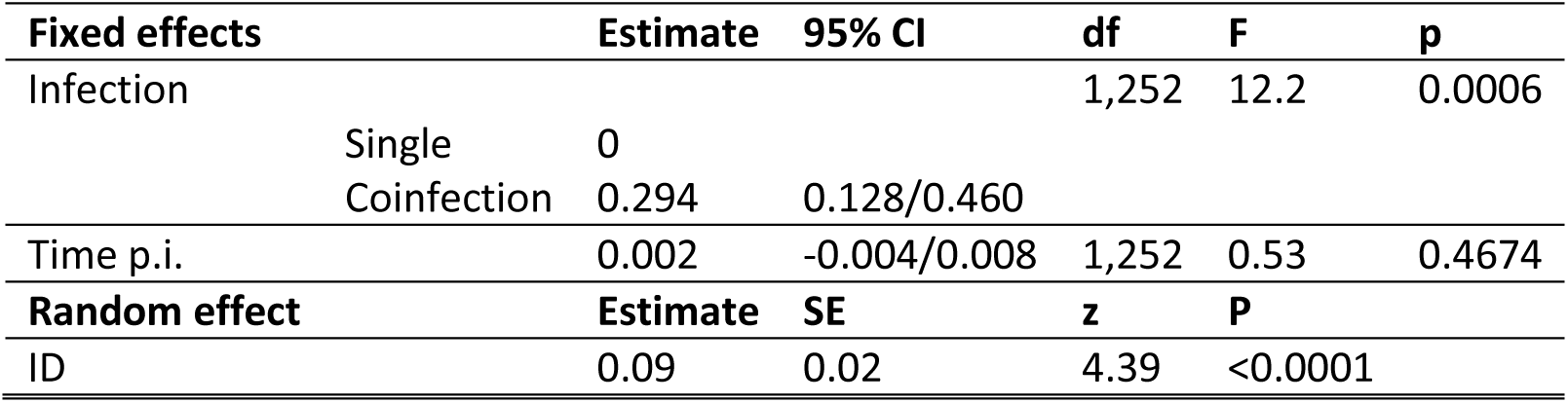
General linear mixed model investigating the changes in fecal egg count (log number of eggs/g of feces) in single Hp infected and coinfected (Py infecting first) mice. We report the parameter estimates with the 95% CI, degrees of freedom (df), F and p values. Mouse ID was included as a random effect to take into account the non-independence of observations for the same individual over time. The Treatment x Time p.i. interaction was not significant and was removed from the model. N = 78 individuals and 331 observations.

| Fixed effects |  | Estimate | 95% CI | df | F | p |
| --- | --- | --- | --- | --- | --- | --- |
| Infection | Single | 0 |  | 1,252 | 12.2 | 0.0006 |
|  | Coinfection | 0.294 | 0.128/0.460 |  |  |  |
| Time p.i. |  | 0.002 | -0.004/0.008 | 1,252 | 0.53 | 0.4674 |
| Random effect |  | Estimate | SE | z | P |  |
| ID |  | 0.09 | 0.02 | 4.39 | <0.0001 |  |

As all mice were infected with the same number of infective larvae and not allowed to be reinfected, this increase in FEC might result from two different non-exclusive processes. First, the environmental conditions provided by a Py infected host might improve the survival of L3 larvae during the prepatent period, resulting in a larger number of adult worms emerging in the intestinal lumen and releasing eggs. Second, more favorable environmental conditions encountered by adult worms in Py infected mice might allow females to produce and release more eggs. Therefore, according to the first hypothesis, the increase in FEC in coinfected hosts should stem from a larger number of adult worms. We compared the biomass of adult worms retrieved at day 21 and 35 p.i. between single Hp infected and coinfected mice and found no difference (when including the coinfection groups separately in the model: GLM, Treatment, F_2,40_ = 0.34, p = 0.7136, Time p.i., F_1,40_ = 0.67, p = 0.4175, fig. S2b; or when clustering them together: GLM, Infection, F_1,41_ = 0.68, p = 0.4132, Time p.i., F_1,41_ = 0.71, p = 0.4037, fig. 1b). Therefore, this result suggests that the increase in egg excretion in coinfected mice is not due to a larger infra-host parasite population size, but rather to an adjustment of worm fecundity.

To go further in the investigation of the process underlying the rise in egg shedding, we assessed whether the infection with Py might induce an increase in FEC when adult worms are already releasing eggs in the intestinal lumen. To this purpose, we infected mice with Hp and after 28 days they were infected with Py and we looked at any change in the number of eggs excreted pre and post Py infection. We found that FEC increased following the Py infection (GLMM, pre vs post Py infection, F_1,91_ = 32.92, p < 0.0001, n = 17 individuals and 109 observations; fig. 1c). As a control, we also compared whether there was a similar increase in FEC post day 28 in single Hp infected mice and found no difference (GLMM, pre vs post day 28 p.i., F_1,82_ = 0.27, p = 0.6049, n = 28 individuals and 111 observations; fig. 1d).

Therefore, overall, these results show that coinfection produced an increase in worm fecundity, and that this is a plastic response that is rapidly induced when the within-host environment changes following the Py infection.

### Immune mechanisms modulating the plastic adjustment of worm fecundity

We then investigated whether the inhibition of the Th2 response due to the infection with Py might account for the observed increase in egg shedding of coinfected mice. We assessed the expression of Th2 cytokine genes in the spleen and found that Hp infection induced an overexpression of both *IL-4* and *IL-13* genes at day 14 and day 21 p.i. compared to non-infected (GLM, *IL-4*, Treatment, F_1,52_ = 346.6, p < 0.0001; Time p.i., F_2,52_ = 0.52, p = 0.5974, fig. 2a, fig. S3a; *IL-13*, Treatment, F_1,49_ = 192.53, p < 0.0001; Time p.i., F_2,49_ = 14.66, p < 0.0001; Treatment x Time p.i., F_2,49_ = 4.83, p = 0.0122, fig. 2b, fig. S3b). However, in agreement with the prediction, both cytokine genes were downregulated in coinfected mice compared to single Hp infected individuals (GLM, *IL-4*, Infection, F_2,99_ = 221.53, p < 0.0001; Time p.i., F_2,99_ = 0.37, p = 0.6883, fig. 2a, fig. S3a; *IL-13*, Infection, F_2,94_ = 84.12, p < 0.0001; Time p.i., F_2,94_ = 25.98, p < 0.0001; Infection x Time p.i., F_4,94_ = 2.73, p = 0.0334, fig. 2b, fig. S3b). When focusing on the coinfection groups separately, the results mirrored those on the FEC, since both cytokine genes were more overexpressed in the Py-14+Hp group, with the Py-28+Hp lying in between the single Hp infected and the other coinfection group (GLM, *IL-4*, Treatment, F_2,71_ = 23.03, p < 0.0001; Time p.i., F_2,71_ = 0.99, p = 0.3750, fig. S3c,e; *IL-13*, Treatment, F_2,70_ = 6.71, p = 0.0022; Time p.i., F_2,70_ = 29.95, p < 0.0001, fig. S3d,f).

**Figure 2.**
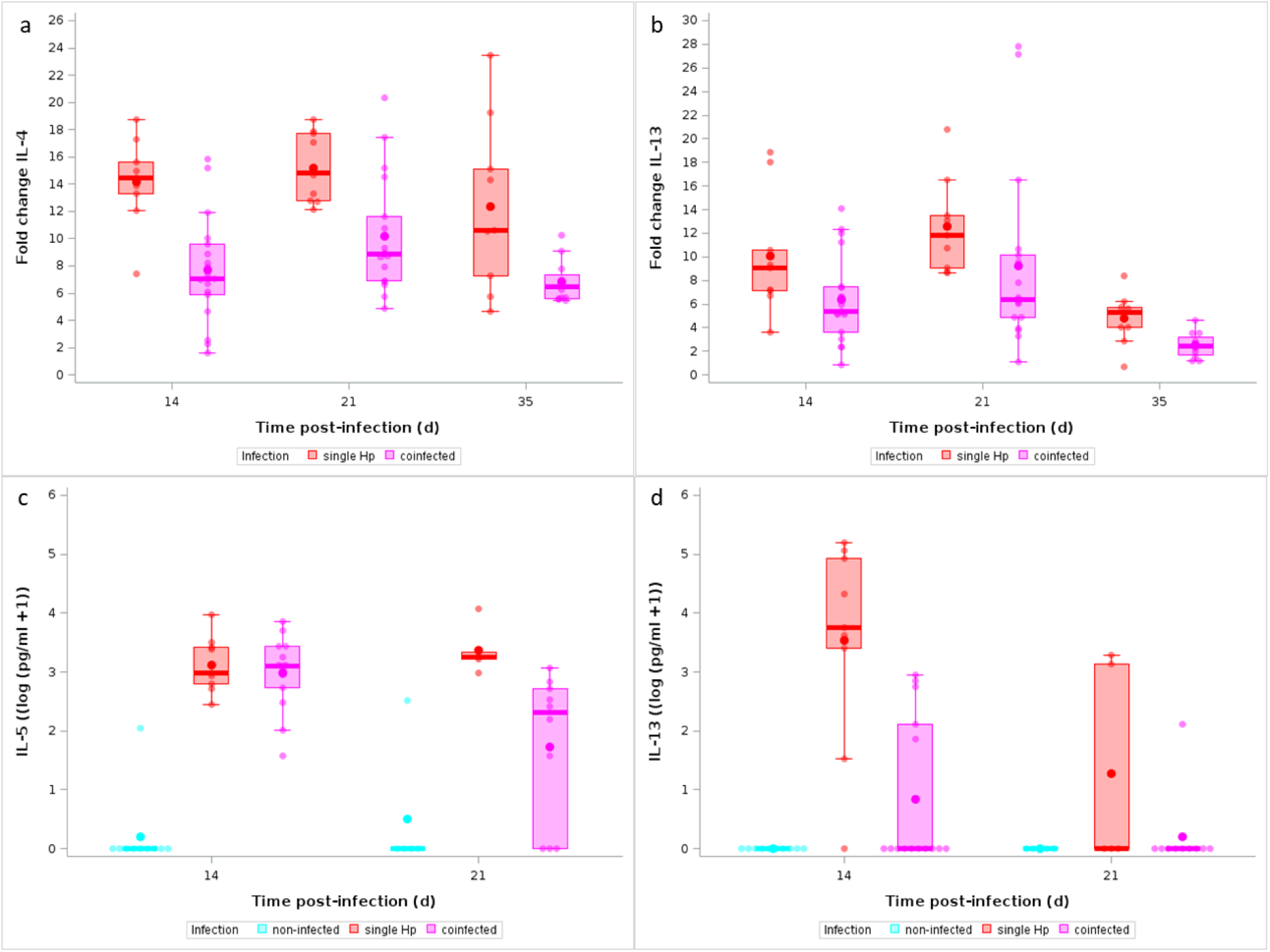
Coinfection with Py inhibits Th2 cytokine production. a) Fold change of IL-4 mRNA in spleen at day 14 and day 21 p.i. in single Hp and coinfected (with Py infecting first) mice; b) Fold change of IL-13 mRNA in spleen at day 14 and day 21 p.i. in single Hp and coinfected (with Py infecting first) mice; c) Levels of IL-5 (pg/ml, log+1) in plasma at day 14 and day 21 p.i. in non-infected, single infected and coinfected (with Py infecting first) mice; d) levels of IL-13 (pg/ml, log+1) in plasma at day 14 and day 21 p.i. in non-infected, single infected and coinfected (with Py infecting first) mice. Dots represent the raw data, the boxes represent the interquartile range (IQR), the horizontal lines the median, and whiskers the range of data within 1.5 the IQR.

We also assessed the amount of circulating IL-5 and IL-13 in plasma and found that Hp infection induced an increase of both cytokines compared to non-infected mice (GLM, IL-5, Treatment, F_1,26_ = 140.63, p < 0.0001, Time p.i., F_1,26_ = 1.11, p = 0.3017, fig. 2c; IL-13, Treatment, F_1,26_ = 31.84, p < 0.0001, Time p.i., F_1,26_ = 4.64, p = 0.0408, fig. 2d). However, confected mice had less amount of IL-5 and IL-13 in plasma compared to single Hp infected (GLM, IL-5, Infection, F_2,48_ = 46.35, p < 0.0001, Time p.i., F_1,48_ = 1.02, p = 0.3167, Infection x Time p.i., F_2,48_ = 5.71, p = 0.0060, fig. 2c; IL-13, Infection, F_2,48_ = 16.66, p < 0.0001, Time p.i., F_1,48_ = 8.17, p = 0.0063, Infection x Time p.i., F_2,48_ = 3.47, p = 0.0393, fig. 2d). Focusing on the coinfection groups separately confirmed the inhibitory effect of coinfection on plasma cytokine levels (GLM, IL-5, Treatment, F_2,33_ = 6.56, p = 0.0040, Time p.i., F_1,33_ = 6.85, p = 0.0133, Treatment x Time p.i., F_2,33_ = 3.43, p = 0.0444, fig. S4a; IL-13, Treatment, F_2,35_ = 10.09, p < 0.0001, Time p.i., F_1,35_ = 6.65, p = 0.0143, fig. S4b).

We finally checked whether coinfection inhibited the expansion of Th2 cells (proportion of GATA-3^+^ within CD4^+^ T cells) in response to Hp infection. The comparison between non-infected and single Hp infected mice confirmed that the infection produced an expansion of the splenic population of Th2 cells over the course of the infection (GLM with a beta distribution of errors, Treatment, F_1,54_ = 209.49, p < 0.0001; Time p.i., F_2,52_ = 6.28, p = 0.0035; fig. 3). However, in agreement with the previous results, coinfection prevented the expansion of Th2 cell population and particularly so at day 14 and 21 p.i., and for the Py-14+Hp group at day 35 p.i. (keeping coinfection groups separated: GLM, Treatment, F_2,69_ = 16.24, p < 0.0001; Time p.i., F_2,69_ = 2.11, p = 0.1295; Treatment x Time p.i., F_4,69_ = 4.26, p = 0.0039, fig. S5; clustering the coinfection groups together: GLM, Infection, F_2,103_ = 100.15, p < 0.0001; Time p.i., F_2,103_ = 4.07, p = 0.0198; fig. 3).

**Figure 3.**
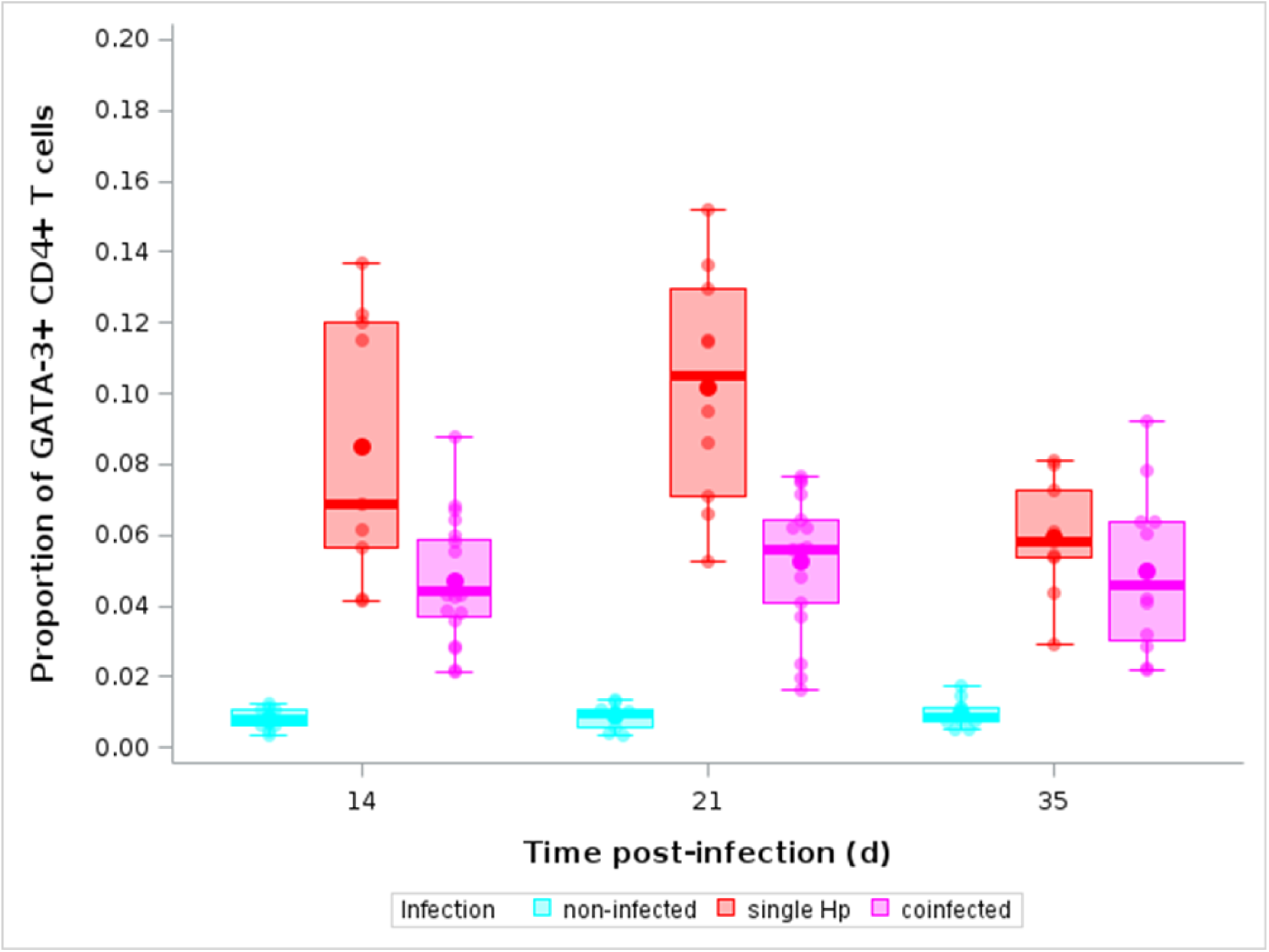
Coinfection with Py represses the expansion of Th2 cells. Proportion of GATA-3+ within CD4+ T cells at day 14, 21 and 35 p.i. for non-infected, single Hp infected and coinfected (with Py infecteing first) mice. Dots represent the raw data, the boxes represent the interquartile range (IQR), the horizontal lines the median, and whiskers the range of data within 1.5 the IQR.

### Manipulation of the Th2 response

If the rise in the number of excreted eggs results from the down regulation of the Th2 response in Py infected mice, we expect that experimentally manipulating the Th2 response in the absence of Py infection should reproduce the results reported above.

To this purpose, we infected mice with Hp and at day 28 p.i., they were treated with neutralizing IL-13 monoclonal antibody, with rIL-13, or with IgG1 as a control, and we assessed whether the treatments produced a shift in egg excretion. For control mice, the injection of IgG1 did not produce any change in egg excretion (GLMM, pre vs post treatment, F_1,64_ = 0.17, p = 0.6857, n = 7 individuals and 72 observations, fig. 4a). However, as expected, the anti-IL-13 antibody treatment produced an increase in egg excretion (GLMM, pre vs post treatment, F_1,68_ = 15.21, p = 0.0002, n = 7 individuals and 76 observations, fig. 4b); while the injection of rIL-13 induced a reduction in egg shedding (GLMM, pre vs post treatment, F_1,79_ = 24.99, p < 0.0001, n = 8 individuals and 88 observations, fig. 4c). Therefore, inhibiting IL-13, which plays a role in the orchestration of the Th2 response, at day 28 post Hp infection, reproduced the results obtained in coinfected mice.

**Figure 4.**
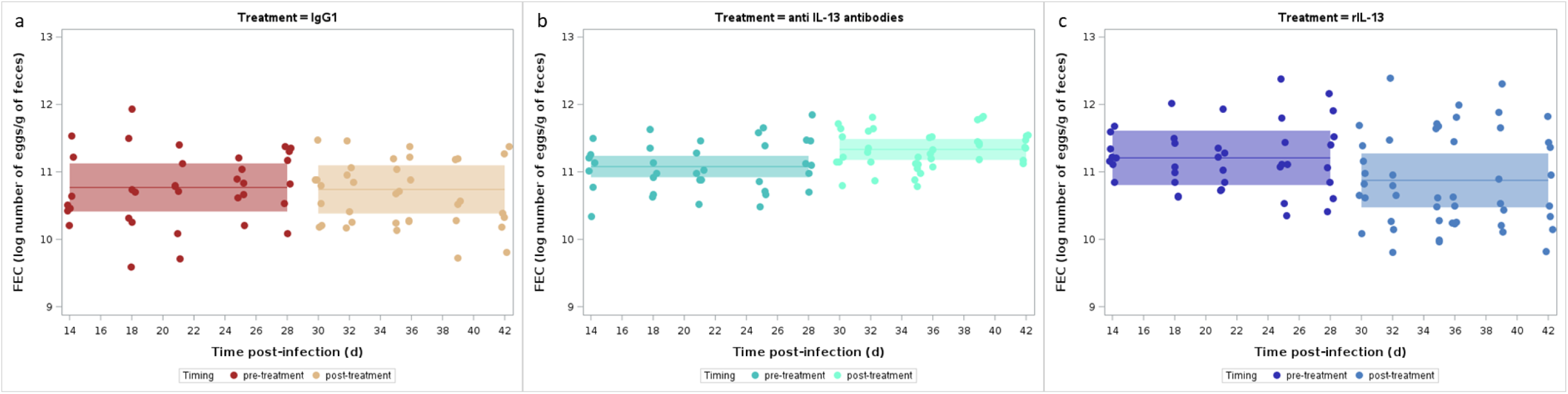
Manipulating Th2 cytokines produces shifts in worm fecundity. a) Changes in FEC pre- and post-injection of IgG1 at day 28, 30 and 32 p.i.; b) Changes in FEC pre- and post-injection of anti-IL-13 antibody at day 28, 30 and 32 p.i.; c) Changes in FEC pre- and post-injection of rIL-13 at day 28, 30 and 32 p.i. Dots represent the raw data, the lines represent the fit of the GLMM, and the shaded area around the line the 95% CI.

### Hp persistence time and relative fitness in single and coinfected hosts

The rise in the number of excreted eggs might be a transitory adjustment to the immune environment provided by hosts coinfected with Py. Moreover, if there is an energetic trade-off between early and late investment in egg production, over the long run, the relative fitness of worms in coinfected hosts should be similar to the fitness of worms in single infected hosts. To investigate this question, we infected and coinfected mice and monitored egg excretion over a 99-day period post Hp infection. Contrary to the prediction, we did not find that the rise in worm fecundity was transitory, since worms in coinfected mice had higher relative fitness compared to worms in single infected hosts (GLM, Treatment, F_2,26_ = 6.14, p = 0.0066, fig. 5a). This result was corroborated by the analysis of the probability of persistence of the infection over time, that showed that single infected mice had a higher probability to clear the infection compared to the coinfection groups (LogRank test, χ²_2_ = 11.01, p = 0.0041, n = 29 individuals, fig. 5b; pairwise comparisons: single Hp vs Py-14+Hp, Sidak adjusted p = 0.0063, single Hp vs Py-28+Hp, Sidak adjusted p = 0.0364, Py-14+Hp vs Py-28+Hp, Sidak adjusted p = 0.8780). Therefore, when Hp infects hosts that had been previously infected with Py, the polarization of the immune response allows the worm to persist for longer and therefore to shed more eggs in the environment.

**Figure 5.**
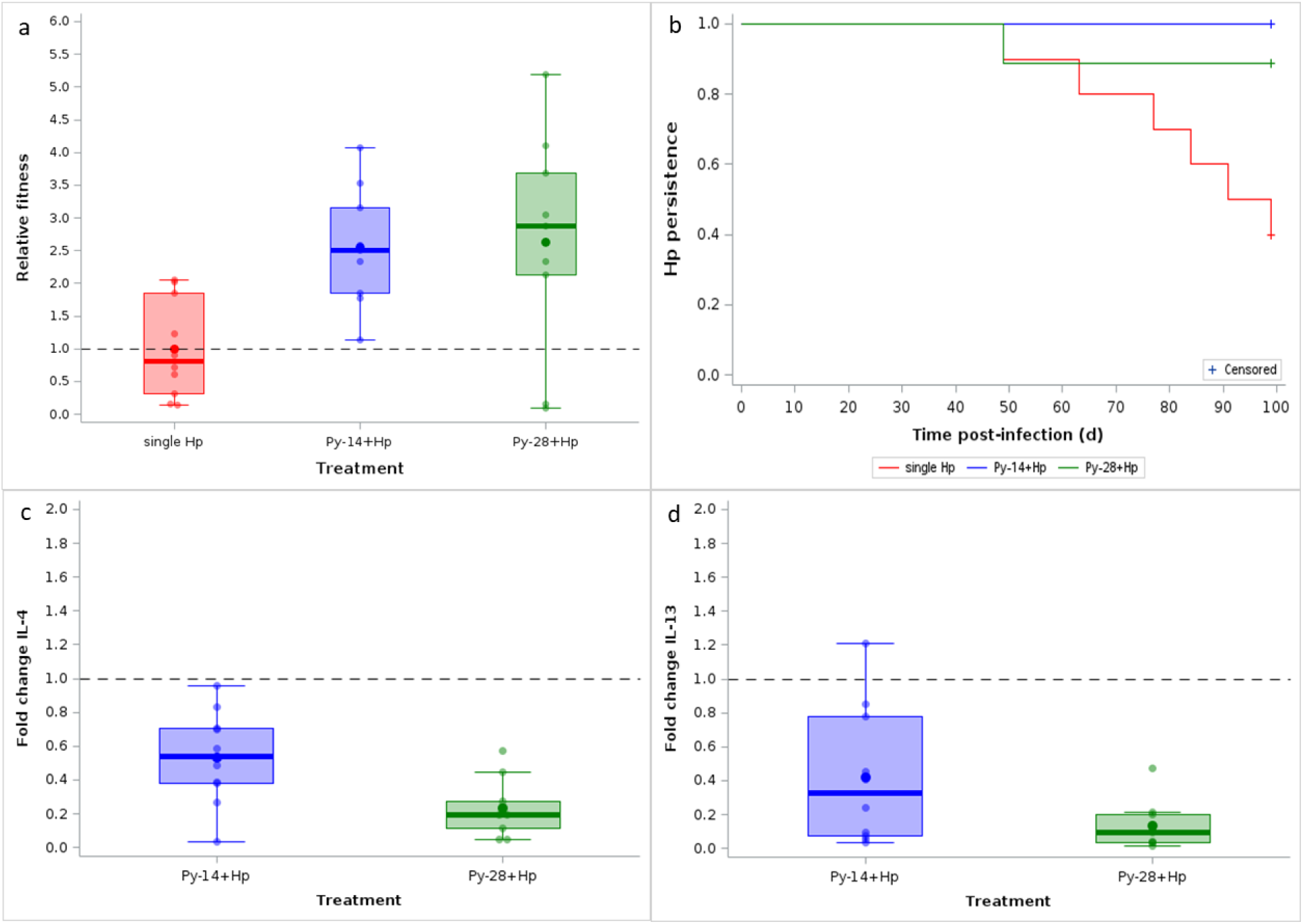
Coinfection with Py improves Hp relative fitness and lengthens the duration of the infection. a) Relative fitness of worms in single and coinfected hosts (with Py infecting first). Dots represent the raw data, the boxes represent the interquartile range (IQR), the horizontal lines the median, and whiskers the range of data within 1.5 the IQR. The dotted line indicates the reference value of single infected hosts. b) Probability of Hp persistence in single and coinfected hosts (with Py infecting first). Crosses indicate censored values. c) Fold change of IL-4 mRNA in spleen at day 99 p.i. in coinfected hosts (Py infecting first and Hp infecting after 14 or 28 days). Dots represent the raw data, the boxes represent the interquartile range (IQR), the horizontal lines the median, and whiskers the range of data within 1.5 the IQR. d) Fold change of IL-13 mRNA in spleen at day 99 p.i. in coinfected hosts (Py infecting first and Hp infecting after 14 or 28 days). Dots represent the raw data, the boxes represent the interquartile range (IQR), the horizontal lines the median, and whiskers the range of data within 1.5 the IQR.

We then investigated whether the down regulation of the Th2 response in coinfected hosts persisted long enough to possibly account for a higher relative fitness of Hp in coinfected hosts. Overall, as expected, both *IL-4* and *IL-13* genes were less expressed at day 99 than at day 14, 21 or 35 post Hp infection, as shown by higher Ct values (GLM: *IL-4*, Time p.i., F_3,101_ = 103.32; p < 0.0001; fig. S6a; *IL-13*, Time p.i., F_3,100_ = 46.64; p < 0.0001, fig. S6b). However, the difference in gene expression between groups persisted at day 99 p.i. (when considering the coinfection groups separately, GLM, *IL-4*, Treatment, F_2,26_ = 9.64, p = 0.0007, fig. 5c, fig. S6c; *IL-13*, Treatment, F_2,26_ = 10.79, p = 0.0004, fig. 5d, fig. S6d; when clustering the coinfection groups together, GLM, *IL-4*, Infection, F_1,27_ = 12.91, p = 0.0013; *IL-13*, Infection, F_1,27_ = 16.04, p = 0.0004).

In the absence of reinfection, the patent period (the period during which hosts keep shedding parasitic eggs) depends on two parameters: the parasite mortality (both immune-dependent and immune-independent mortality), and host mortality. In a previous work, we showed that coinfection between Hp and Py increases the severity of the disease in an asymmetrical manner, the disease becoming more severe only when Py infects hosts already harboring Hp (Dusuel et al. 2025). We therefore investigated whether host mortality that occurs in coinfected hosts when Py follows Hp, might offset any potential benefit in terms of more favorable immune environment and therefore results in a decreased lifetime reproductive success. To explore this question, we first compared the persistence of the Hp infection in single and coinfected mice (when Py infects mice that had been harboring Hp during the previous 28 days) and indeed found that the two groups had similar persistence time, although for different reasons (parasite mortality in the single infection group and host mortality in the coinfection group) (LogRank test, χ²_1_ = 0.042, p = 0.8369, n = 19, fig. 6a). However, despite similar infection persistence time, Hp in coinfected hosts had substantially higher lifetime reproductive success compared to Hp in single infected hosts (GLM, F_1,17_ = 31.83, p < 0.0001, fig. 6b). Therefore, coinfection with Py provided a net fitness benefit to Hp, even in a coinfection group where host mortality contributes to stop Hp transmission.

**Figure 6.**
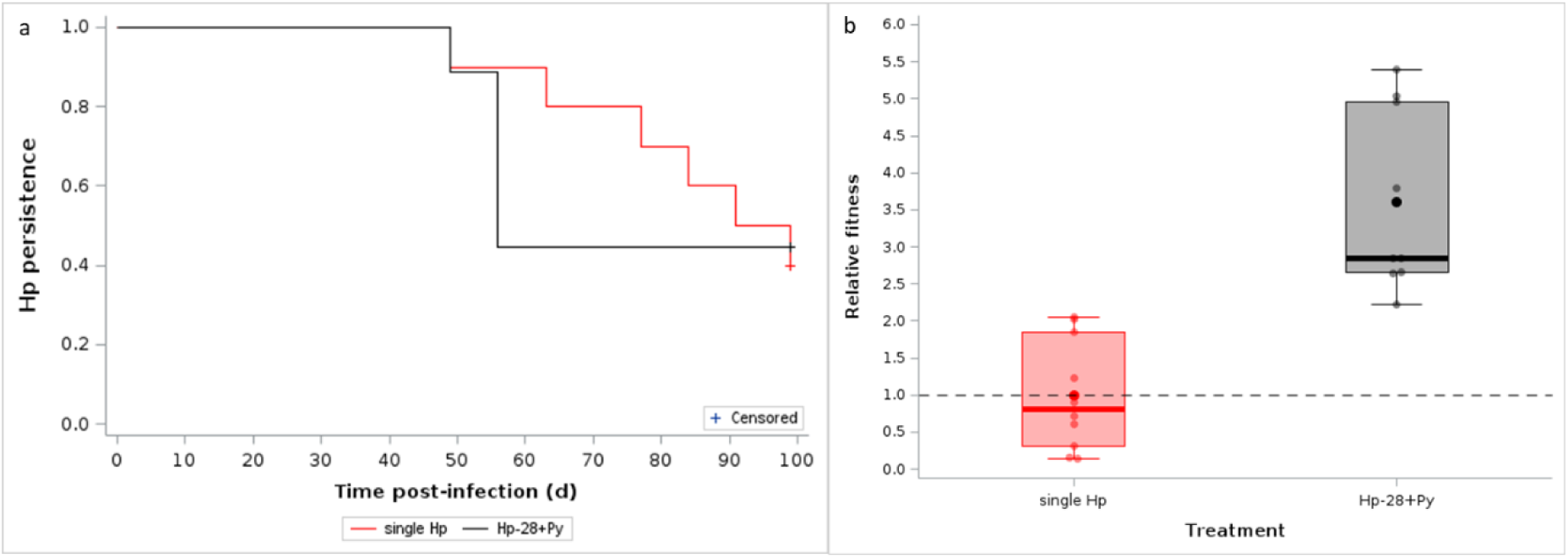
Coinfection with Py improves Hp relative fitness even when host mortality stops egg shedding. a) Probability of Hp persistence in single and coinfected hosts (with Hp infecting first). Crosses indicate censored values. b) Relative fitness of worms in single and coinfected hosts (with Hp infecting first). Dots represent the raw data, the boxes represent the interquartile range (IQR), the horizontal lines the median, and whiskers the range of data within 1.5 the IQR.

## Discussion

We found that coinfection with a malaria parasite has profound effects on the dynamics and fitness of a gastrointestinal nematode. We further show that these effects are plastic, long lasting responses, mediated by changes in host immunity. Therefore, coinfection with *Plasmodium* has the potential to profoundly alter the epidemiology of STHs.

Micro and macroparasites elicit different facets of the host immune system that have reciprocal inhibitory effects (Paludan 1998, Patel et al. 2009, Ezenwa and Jolles 2011, Coomes et al. 2015, Kapse et al. 2022, Seguel et al. 2022). Therefore, hosts harboring a microparasite infection should offer a more favorable ground for the infection with helminths, both in terms of the likelihood of establishing the infection and in terms of production of parasite propagules. For instance, mice infected with the gastrointestinal nematode *Nippostrongylus brasiliensis* and coinfected with the protozoan *Toxoplasma gondii* had lower Th2 responses as assessed by Th2 cytokine production in the mesenteric lymph nodes and the spleen, and higher and prolonged nematode egg excretion

(Liesenfeld et al. 2004). In a slightly different system of concomitant infection with *Nippostrongylus brasiliensis* and *Plasmodium chabaudi*, Hoeve et al. (2009) also reported an impaired Th2 response. Similar results were also reported in mice coinfected with *Plasmodium chabaudi* and Hp (Coomes et al. 2015). In agreement with these previous results, we found that Py infection weakened the Th2 response and this correlated with higher parasite fecundity and longer patent period. On the contrary, we did not find strong evidence suggesting that previous infection with Py improves the infectivity of larvae, defined as the probability of molting into reproductive adults, since the dry mass of adult worms retrieved in the intestinal lumen did not differ between single infected and coinfected mice. Asymmetrical effects of altered Th2 response due to coinfection with microparasites have already been reported in similar systems. For instance, mice infected with Hp and subsequently infected with *Leishamia infantum* (7 days post Hp infection) resulted in higher Hp egg shedding, while the larval infectivity was not affected by the coinfection (González-Sánchez et al. 2018). Enhanced Hp fecundity, independent from larval survival, has also been found in mice that had been previously infected with *Toxoplasma gondii* (Ahmed et al. 2017). In all these examples, including our study, the immune-driven effect of coinfection therefore involved plastic adjustments of reproductive effort, rather than an overall improvement of infectivity. In support to this hypothesis, we also provided additional evidence indicating that coinfection had a direct effect on worm fecundity. Indeed, when mice harboring a chronic infection with Hp were subsequently infected with Py they excreted a higher number of nematode eggs in the feces. This result unambiguously shows that worms were able to rapidly adjust their investment into egg production, because at day 28 post Hp infection all larvae have molted into adult worms and in the absence of reinfection, the population size of reproductive worms cannot increase. Whatever the demographic parameters affected by the coinfection, these results also show that the increase in egg shedding does not depend on the order of infection, since we found that mice that were either infected first with Py or with Hp excreted more nematode eggs compared to single infected hosts. We believe that this finding has relevant consequences for the understanding of the epidemiology of STHs. Indeed, order of infection has been suggested to be an important driver of the infection dynamics in a few model systems (Karvonen et al. 2019). This raises the question of the importance of the order of infection under natural conditions where hosts are exposed to multiple pathogens and parasites that have different life cycles and different host age-dependent probability of infection. However, in the current case, the effect of malaria infection on Hp egg shedding appears to be independent of infection order. Therefore, it is straightforward to predict that, admitting that these results can be extrapolated to other helminth/malaria species, coinfection should have a consistent effect on the spreading and the transmission of gastrointestinal nematodes. Indeed, for STHs that have simple life cycle and do not rely on intermediate hosts to complete their cycle a key epidemiological parameter is the quantity of eggs/infective larvae released in the environment, since transmission occurs when susceptible hosts get into contact with the transmissible stages of the parasite. Therefore, everything else being equal, the higher the reservoir of transmissible stages in the environment, the higher the probability of successful transmission. Another consequence of these results is that treatments aiming at reducing the risk of coinfection might also directly jeopardize the transmission success of soil-transmitted nematodes.

One of the central tenets of life history theory is that when resources are limited, increased investment into reproductive effort at a given stage (age) should result in decreased investment into reproductive effort at later stages (ages) or increased mortality rate (Stearns 1992). Based on this reasoning, we predicted that the enhanced egg shedding that we observed from day 11 to day 35 p.i. should be paid later on, finally resulting in similar fitness of worms in single infected and coinfected mice. Contrary to this prediction, we found that worms in coinfected mice had consistently higher fitness (cumulative number of excreted eggs) relative to worms in single infected hosts. In particular, we showed that worms in coinfected hosts had longer persistence time and therefore a longer patent period allowing them to cumulatively shedding more eggs in the environment. The absence of trade-off and the finding of higher relative fitness of worms in coinfected hosts suggests that the host immune response strongly constraints the expression of the demographic traits of Hp (survival and fecundity) and that relaxing these constraints (e.g., in coinfected hosts) allows the nematode to release more propagules over time.

For directly transmitted parasites (i.e., parasites that do not rely on vectors or intermediate hosts), fitness (R_0_) positively depends on i) the probability of encounter with susceptible hosts, ii) the probability of successful inter-host transmission and iii) the duration of the infectious period (the period during which the parasite keeps producing transmissible stages) (Earn 2008). Therefore, everything else being equal, the longer the duration of the infectious period, the better the fitness. In the absence of reinfection, the persistence of the infection depends on the survival of the worms that includes the background mortality and the immune-mediated mortality, and the survival of the host. Therefore, when coinfection produces a higher risk of host mortality, parasite fitness might be jeopardized. We previously showed that when hosts are first infected with Py and subsequently with Hp, the disease symptoms are similar to the single infection, with no host mortality (Dusuel et al. 2025). Therefore, the difference in the persistence of the Hp infection in single infected and coinfected mice only stems from differential adult worm survival (both immune-dependent and immune-independent mortality). However, when Py infects hosts that already harbor Hp, the risk of host mortality is higher than for single infection, contributing to shorten the patent period (Dusuel et al. 2025). In agreement with this, we found that the persistence of the Hp infection did not differ between single infected and coinfected hosts where Hp infects first, although the underlying reasons of this similar persistence were different: worm mortality vs. host mortality. However, despite similar duration of the infection, worms in the coinfection group still had a much higher relative fitness, showing that the increase in fecundity associated with a favorable immune environment largely outweighs the cost of host mortality. It is important to acknowledge that our assessment of Hp demographic and fitness traits do not refer to individual worms but to the whole infra-host nematode population. Indeed, for obvious reasons, it is not possible to monitor individual worm fecundity and survival. Moreover, we only used a proxy of overall egg excretion since we measured FEC on a weekly basis. Finally, our conclusion of better worm fitness in coinfected hosts rests on the assumption that higher investment into egg production does not alter other key life history traits of the parasite, such as hatching success, larval survival in the external environment or larval infectivity.

We also showed that the differences in the life history traits of worms in single infected and coinfected mice are likely to be driven by the immune environment provided by the host. In agreement with previous results, we found that Hp infection induced a Th2 response characterized by the upregulation of Th2 cytokine genes, higher levels of circulating Th2 cytokines and an expansion of Th2 cells (GATA-3^+^ CD4^+^ T cells) in the spleen. However, coinfection consistently weakened the anti-Hp immune response, since coinfected hosts had downregulated Th2 cytokine gene expression, reduced levels of circulating Th2 cytokines and reduced expansion of Th2 cells. We interpreted these inhibitory effects as the consequence of the polarization of the immune response towards anti-Py effectors (Th1), known to have inhibitory effects on the Th2 response (e.g., Ahmed et al. 2017). Alternatively, condition-dependent effects might also contribute to explain the rise in Hp fecundity, if coinfected hosts suffer from worst body condition. To disentangle these two hypotheses, we conducted an experiment where we manipulated the Th2 response up and down, in the absence of Py infection. Therefore, if the rise in Hp fecundity is mediated by immune effects, we should expect that inhibiting or enhancing the Th2 response, in the absence of Py infection, should reproduce the results observed in coinfected hosts. Indeed, we found that treating Hp infected mice with an anti-IL-13 monoclonal antibody increased egg excretion, while treatment with an rIL-13 reduced egg shedding. Therefore, these results further show that the adjustment of fecundity does not reflect a parasite response to poor host condition, caused by the coinfection, and corroborate previous work showing that IL-13 and its receptor are important component of the anti-Hp response (Sun et al. 2016). Manipulation of the Th2 or the Th1 response using a variety of approaches has provided consistent results, similar to those reported here. For instance, blockade of IFN-γ with anti-IFN-γ antibody restored the Th2 response in a coinfection model between Hp and *Plasmodium chabaudi* (Coomes et al. 2015), while supplementing mice with IFN-γ increased Hp fecundity (Affinass et al. 2018). Blockade or supplementation with IL-4 also had the expected results on Hp fecundity and infection persistence (Urban et al. 1991, 1995).

Manipulation of IL-4 had no effect on the larval stage (Urban et al. 1995) which is also in agreement with the reports of no effect of coinfection on larval survival. IL-4 and IL-13 have also been shown to play a role in the expulsion of adult worms by promoting peristaltic movements and the contractility of intestinal smooth muscles (Zhao et al. 2003).

Interestingly, we found that the difference between single infected and coinfected hosts in the expression of Th2 cytokine genes persisted well after the Py infection had been cleared. Of course, the amount of *IL-4* and *IL-13* mRNA was lower at day 99 p.i. compared to day 14 or day 21 p.i., but the expression of both genes were still downregulated in coinfected hosts. Although a long lasting anti-Hp Th2 response has already been reported up to day 70 p.i. (Finney et al. 2007), we cannot affirm that the longer persistence and higher fitness of worms in coinfected hosts is due to a permanent downregulation of the Th2 response in coinfected hosts. Further work should elucidate this hypothesis and the possible mechanisms underlying a prolonged inhibition (once Py has been cleared) of the Th2 response in coinfected hosts.

To conclude, these results shed light on the possible effect of malaria coinfection (and more generally of microparasite coinfection) on the egg excretion and within-host persistence of a gastrointestinal nematode. In particular, they show that coinfection affects some key epidemiological and demographic traits, generating positive feedback whereby coinfected hosts contribute to maintaining a large reservoir of transmissible stages in the environment, which in turn sustains a high risk of coinfection. Assuming these results can be extrapolated to other systems (involving human microparasites and STHs), they suggest that targeting the coinfecting microparasites, in addition to the direct benefit for the host, might also indirectly reduce helminth transmission success by reducing the number of transmissible stages found in the external environment, and as such reduce the risk of infection.

## Acknowledgments

We are grateful to Valérie Saint Giorgio and all the staff of the animal facility for taking care of the animals.

## Author contribution

GS, BF, MR and BR conceived the study and gathered the funding; AD, LB, GS and BF collected the data; AD, LB and EG performed the lab work; GS analyzed the data; GS, AD and LB wrote the first draft of the manuscript; all authors revised and approved the final version.

